# Experimental manipulation of nocturnal nest cavity temperature in wild blue tits

**DOI:** 10.1101/279455

**Authors:** Emily G. Simmonds, Ben C. Sheldon, Tim Coulson, Ella F. Cole

**Affiliations:** Department of Zoology, University of Oxford, Oxford, UK; Department of Mathematical Sciences and Centre for Biodiversity Dynamics, Norwegian University of Science and Technology (NTNU), Trondheim, Norway.

**Keywords:** blue tits (cyanistes caeruleus), phenology, experimental manipulation, lay date, in-nest temperature, heating and cooling

## Abstract

Advances in the timing of reproduction in temperate species are some of the most well documented biotic responses to increasing global temperatures. However, the magnitude and rate of these advances in timing are not equal across all taxonomic groups. These differences can lead to disruption of interspecific relationships if species respond differently to temperature changes. Understanding the relationship between temperature and phenology is a key step in predicting future population trends for species living in seasonal environments. However, experimentally manipulating temperature in the wild is logistically challenging and has consequently rarely been attempted. In this study we experimentally test whether in-nest temperatures in early spring act as a cue for breeding phenology in a population of wild blue tits (*Cyanistes caeruleus*). We split nests into three treatments; heated, cooled, and control. In-nest temperature in the heated and cooled boxes was manipulated by an average of ± 0.6 °C from control temperatures using heating devices and ice packs respectively. We assessed the impact of our experimental manipulation on box occupancy and reproductive timing. We found trends towards earlier phenology in heated nest boxes in addition to a higher occupancy rate in cooled boxes, however neither of these trends was found to be statistically significant. Our ability to distinguish statistical signals was hampered by unexpectedly low occupancy rates across all experimental treatments. Based on the results we cannot say if nocturnal in-nest temperature is an important cue for nest box choice or the timing of laying.

## Introduction

Advances in phenology, the timing of life history events, have been widely documented amongst temperate species over recent decades, with examples spanning multiple ecosystems and almost all taxonomic groups (Thackeray et al. 2010). These advances have coincided with increases in global temperatures as a result of anthropogenic climate change (Menzel et al. 2006; Parmesan 2006; Singer & Parmesan 2010). Failure to time breeding to match with the peak abundance of resources can result in energetic demands that cannot be met and a consequent reduction in fitness (Reed, Grøtan, et al. 2013; Reed, Jenouvrier, et al. 2013). Therefore understanding how temperature influences breeding phenology is crucial for predicting the impact of climatic change on reproductive success. Despite this importance, the precise role that temperature plays in altering breeding phenology has not yet been determined.

Matching peak energetic demands to a shifting resource peak is particularly challenging for consumer species, which rely on species in lower trophic levels as their resource. The consumer species must respond to temperature in the same way as the resource in order to maintain synchrony, this is challenging for species with temperature independent development, where embryos have a fixed developmental period. In contrast to species with temperature-dependent development, which respond directly to temperature changes (Perrins 1979), temperature-independent species must use predictive cues in order to determine the optimal breeding time (van Noordwijk et al. 1995; Perrins 1979). Not all species respond to the environment in the same way, this can have significant impacts on interspecific interactions (Thackeray et al. 2010; Parmesan 2007; Visser & Holleman 2001; Plard et al. 2014). If interacting species differ in their response to environmental cues or the specific cues they use to time their phenology, they will shift their timing to differing degrees, which can disrupt synchrony between species (Cushing 1969). For instance, if a prey species advances their phenology at a faster rate than a predator species, due to greater responsiveness to an environmental cue this could result in temporal mismatch. Peak abundance of resources would occur prior to the peak energetic demand, reducing survival and reproductive success of the predator (Reed, Grøtan, et al. 2013). This has been the case for great tits (*Parus major*) in the Netherlands. Great tits in the Hoge Veluwe population have not advanced their phenology as much as resources despite increases in spring temperatures, whereas peak abundance of their prey species, winter moth caterpillars (*Operapthera brumata*) has advanced by over one week in two decades (Visser et al. 1998). This has resulted in fitness reductions for these great tits (Reed, Grøtan, et al. 2013). In order to understand how and where these mismatches may occur, it is important to identify the precise cues that individuals use to time their reproductive behaviour. Only through understanding the mechanisms that control reproductive timing can we predict how timing will change into the future and whether temporal mismatch will be likely.

Attempts to identify the cues that drive reproductive phenology consist of several approaches. The most commonly used approach is to conduct regression based analyses on observational data (van Noordwijk et al. 1995; Charmantier et al. 2008; Menzel et al. 2006; Parmesan & Yohe 2003; Torti & Dunn 2005). This approach typically consists of regressing candidate cues (often temperature) against a phenological event of interest. Either testing different temporal windows or days of temperature (Roberts 2008; Thackeray et al. 2016; van de Pol et al. 2016; Phillimore et al. 2013). The results of studies of this type, have shown that the breeding phenology of northern temperate species is highly correlated with various measures of spring temperature (Visser et al. 1998; van Noordwijk et al. 1995; Charmantier et al. 2008; Cleland et al. 2007; Menzel et al. 2006; Parmesan & Yohe 2003). April temperatures were shown to correlate with clutch initiation date in gulls (Brommer et al. 2008) and temperatures during March and April explained 70 % of the variation in clutch initiation date for great tits and blue tits (*Cyanistes caeruleus*) in the UK (Perrins & McCleery 1989). However, it can be difficult to pinpoint exact temperature cues through these analyses given that temporal windows and the temperature variables tested are all at least partially correlated. Furthermore, regression based studies do not allow the determination of causality, they can only assess if two variables correlate. In order to establish causal links between environmental cues and phenological events we must therefore use experimental manipulations are required.

Experimental manipulations of temperature cues pose a logistical challenge, particularly for large vertebrate species. While sessile species, such as plants, can be manipulated in situ, mobile species such as most animals, cannot. Animal species tend to cover considerable distances, daily, weekly or seasonally, experiencing temperature cues across a large spatial and temporal range. Consequently, manipulating temperature across these scales in the wild is often impossible. As a result, captive experiments provide one alternative (Schaper et al. 2011; Schaper et al. 2012). These captive experiments provide a good environment to determine the role of different cues and test causality. For instance, captive experiments on great tits used temperature controlled aviaries to explore the influence of temperature on clutch initiation date. This experiment showed that while an increasing temperature caused clutch initiation date to advance, changes to mean temperature or temperature variation alone did not impact phenology (Schaper et al. 2012). However, experiments in captivity do not tell us whether these relationships would hold in the wild. There have been examples of experimental manipulations failing to capture observed patterns (Lambrechts et al. 1999; Wolkovich et al. 2012). Captive environments have highly controlled conditions, often investigating a single focal cue at a time (Schaper et al. 2011; Schaper et al. 2012) and consequently failing to take account of the influence of other variables. It is unlikely in reality that a single driver determines all phenological change. Furthermore, captive environments create artificial conditions, which are potentially stressful for the animals and can lead to adjustment of behaviour away from what it would be in the wild (Lambrechts et al. 1999).

To identify whether relationships identified during captive experiments or regression analyses also hold in the wild we need to perform experiments on natural systems. Cavity nesting birds, such as great tits and blue tits, provide a good study system for such experiments. Reproductive timing plays an important role in reproductive success in these species, as a failure to time the peak energetic requirement of reproduction with the peak availability of food resources can result in reduced reproductive success (Reed, Grøtan, et al. 2013). There have also been many statistical assessments of cues on these populations (Husby et al. 2010; van Noordwijk et al. 1995; Perrins & McCleery 1989; Perrins 1965b) and some captive experiments (Schaper et al. 2012), which can be used to inform field experiments. Additionally nest cavities provide an enclosed environment that can be manipulated. Previous studies that have experimentally heated and cooled nest cavities, have shown that cavity temperature influences nocturnal incubation (Vedder 2012) and the incubation intensity (Ardia et al. 2010) in blue tits and tree swallows (*Tachycineta bicolor*), respectively. In contrast, in spite of observational evidence that suggests nest temperatures influence clutch initiation date (Dhondt & Eyckerman 1979; O’Connor 1978) similar experimental manipulations have been unable to demonstrate this effect. A previous experiment on great tits attempting to manipulate clutch initiation date (Nager & van Noordwijk 1992), through heating and cooling nest boxes in two consecutive years, failed to identify a significant signal of temperature on clutch initiation date. This experiment heated and cooled groups (N = 40 boxes per year) of nest boxes at night by ± 0.75 °C from two to four weeks prior to the median clutch initiation date of each year; however, no difference in phenology was detected between treatments. Here we repeat and extend this study using a population of wild blue tits. We heated and cooled 87 (N = 29 per treatment) nest boxes by an average of ± 0.6 °C from 6 pm to midnight from 17 days before the first egg was laid, and testing whether this manipulation influences nest box choice, commencement of nest building and clutch initiation date. We hypothesise that female blue tits will preferentially choose to nest in warmer nest boxes in order to minimise their nocturnal energy expenditure (Ardia et al. 2009; Vedder 2012; Dhondt & Eyckerman 1979). Furthermore, we predict that the temperature of the nest cavity will also act as a cue for clutch initiation date (Dhondt & Eyckerman 1979; O’Connor 1978), therefore the warmer nests will also experience the earliest clutch initiation dates and the coolest boxes, the latest clutch initiation date.

## Methodology

### The Study system

This study was conducted in spring 2015 using a population of wild blue tits in Wytham Woods, UK. The nest box breeding population of blue tits has been studied in detail using standardised procedures (Perrins 1965a; McCleery & Perrins 1985) since 2002. Annually 250-450 pairs of blue tits breed in the woodland (Perrins 1979). 147 blue tit nest boxes were erected across the woodland in 2002, with a further 41 boxes added in 2008. These blue tit boxes have entrance holes with a diameter too small to allow larger species, such as great tits, to enter the boxes. By using these boxes rather than those with larger entrance holes, which are also present in the study area, we were able to exclude great tits from our experimental manipulations. We use 87 of these blue tit nest boxes as the focus of our study (Figure 1). These 87 were chosen because they are located close together in a topographically similar area of the woodland, consequently limiting habitat differences. Proximity of the boxes was essential for this experiment to ensure all experimental treatments could be set running within a two hour window each evening.

**Figure 1:**
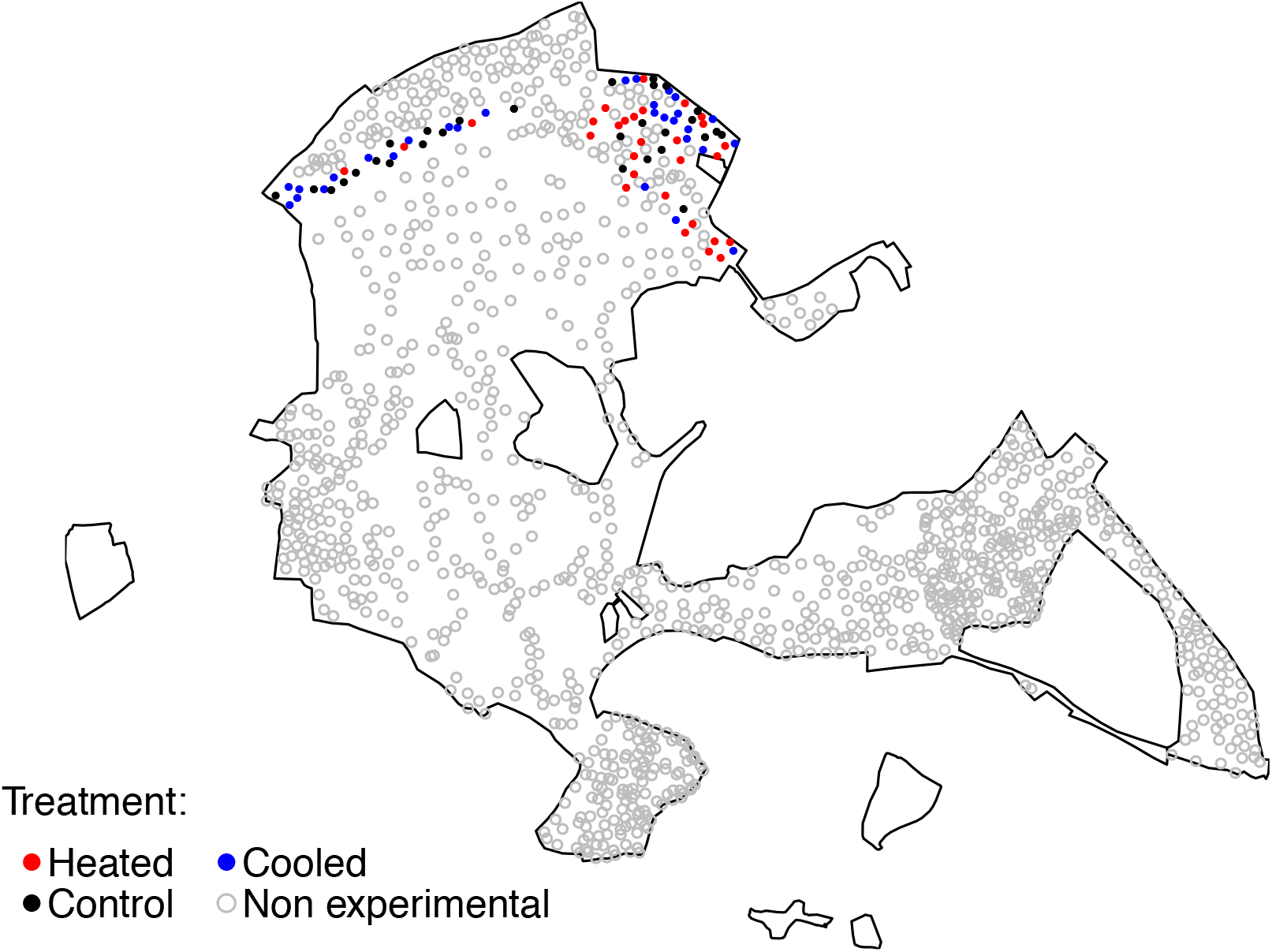
Map of position of experimental (filled circles) and non-experimental (open circles) nest boxes in Wytham woods. Experimental treatments are colour coded.

### Environmental data collection

Altitude data were extracted by Wilkin et al (Wilkin et al. 2006) from an Inverse Distance Weighting interpolation of a 50 m resolution Land Form profile Digital-Terrain-Model data set provided by Ordnance Survey.

Local ambient temperature was collected via a grid of ambient temperature iButtons (DS1923-F5, accurate to ± 0.5 °C, HomeChip Ltd) set to measure absolute temperature every 30 minutes. 200 of these ambient temperature iButtons were distributed in a grid system across Wytham woods with positions chosen to reflect the density of nest boxes. QGIS (Team 2016) was used to match each experimental nest box to a nearest ambient temperature iButton, therefore allowing the calculation of local ambient temperature for each box. The measure of local ambient temperature used in this study is mean temperature for the experimental period, from 1^st^ April to 6^th^ May or until the first egg is laid, whichever is first.

Long-term temperature data is taken from the Radcliffe Meteorological Station (located 5 km East of Wytham Woods). This station records absolute temperature during the day and calculates daily minimum and maximum temperatures from 1815 to the present day (Radcliffe Meterological Station n.d.).

### Experimental Manipulation and Phenological Data Collection

Of the 87 experimental boxes, 29 were assigned to each of the three treatment types: heated, cooled, and control. Boxes were assigned to treatments using a stratified sampling technique, taking account of historical box popularity. Historical popularity was determined by calculating the proportion of years in which each box has been occupied since its placement in the woodland. Boxes were then grouped into bands of popularity in increments of 10 %, within each group treatments were assigned randomly using R (Team 2008). This assignment method was used to prevent a clustering of treatments by historical popularity as this metric showed a significant negative correlation with mean clutch initiation date of the nest box (EST = −3.18, SE = 1.46, P = 0.03), with the historically most popular boxes having the earliest clutch initiation dates. This sampling approach also created an even likelihood of box occupancy across treatments, therefore allowing us to explicitly test whether our experimental manipulation altered the popularity of individual boxes. We tested whether other spatial components, such as altitude, influenced the mean clutch initiation date of each box, but this was not statistically significant (EST = 0.02, SE = 0.03, P = 0.52). Consequently, we did not take account of altitude when assigning treatments. This assignment of treatments does mean individuals can move between different treatments if they do not roost in the same nest box every night, however, it also avoids temporal or spatial clustering of treatments and is essential for the exploration of in-nest temperature influence on occupancy.

Manipulations began on the 27^th^ of March 2015, 17 days before the first egg was laid in the experimental boxes. Manipulations occurred every evening, ending when the first egg was laid in a box or on the 6^th^ May 2015 for unoccupied boxes, 30 days after the first egg was laid in the whole woodland. The cut off of 30 days after the first egg is a common measure used to exclude second clutches and repeat clutches which result from early failures (Van Der Jeugd & McCleery 2002). The heating device comprised (see Figure 2 for details of device construction) of a three-volt filament bulb secured in a mount within a metal topped polystyrene disc (approximately 2.5-3.5 cm thick), which was placed on top of a plastic ring with a 5 cm gap at the front (also 2.5 cm in height). The bulb was powered by two rechargeable batteries, size C, located beneath the polystyrene disc within the plastic ring (see Figure 2). The bulb was covered from above by a 5 cm diameter metal disc, to hide the light produced. Heating was achieved by the bulb heating the metal disc and convectional transfer of the heat up into the nest cavity. These heated boxes also contained four, non-frozen, Thermafreeze ice packs (Thermafreeze EU, black ice packs) in the same configuration as the cooling device (see below) to ensure that all experimental boxes had the same outward appearance.

**Figure 2:**
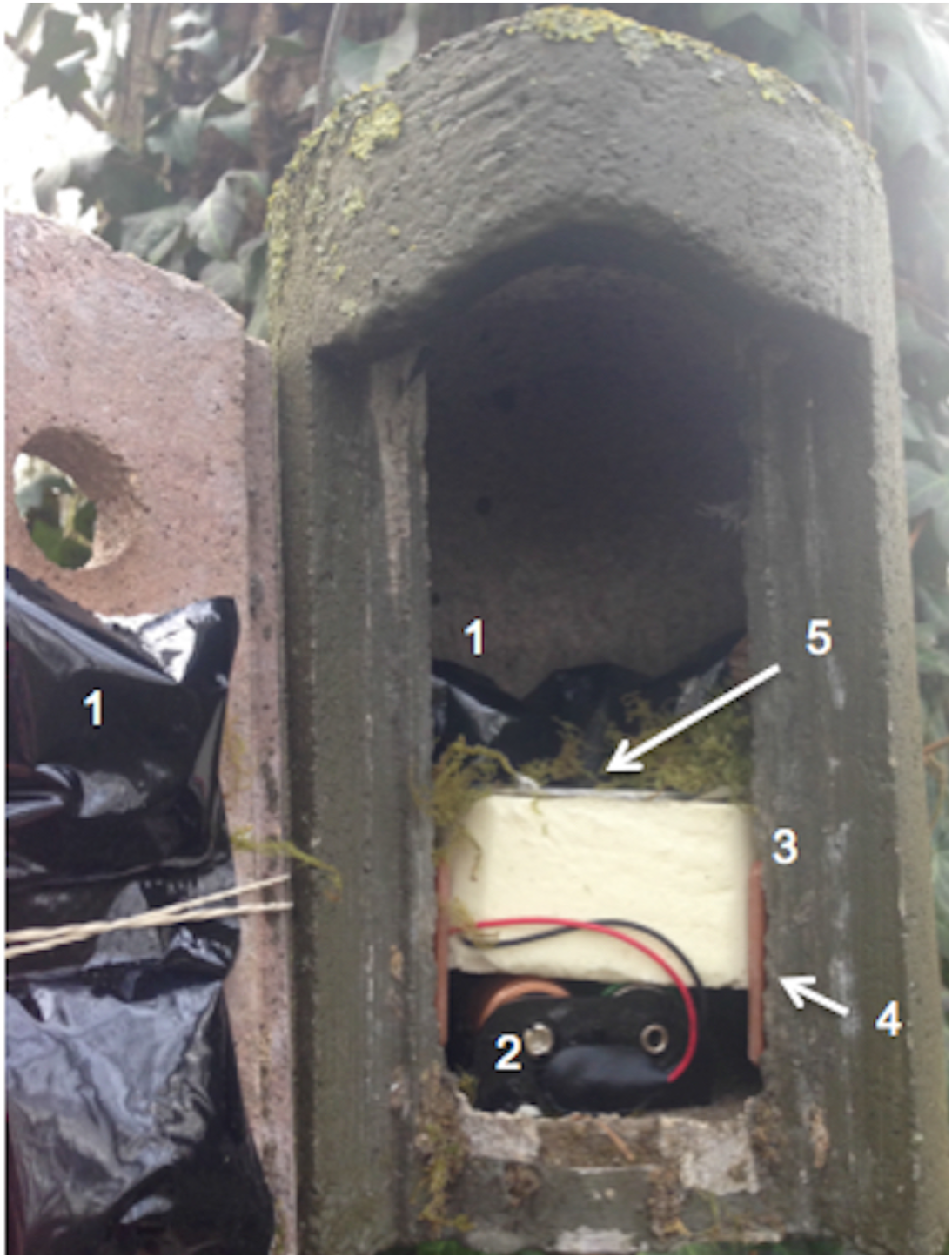
Photograph of experimental set up for a heated box. Key components of the set up are numbered 1) Ice packs mounted front and back of the box, 2) battery pack, 3) polystyrene insert, 4) plastic ring to hold polystyrene above the battery pack, 5) metal disc topping the bulb. In control and cooled boxes, the battery pack would be replaced by a second polystyrene insert.

The cooling device comprised two frozen Thermafreeze ice packs secured at the back of each nest box and two on the nest box door. Ice packs at the back of the box were secured using the same metal topped polystyrene disc as used in the heating device, with another polystyrene disc below (around 2.5 cm thick) to achieve the same thickness as the heating device but without the plastic ring and batteries. Ice packs on the nest box door were secured using rubber bands. All ice packs had a black outer side to camouflage them from the birds once the nest box was closed.

The control treatments comprised the same construction as the cooled treatment but with non-frozen ice packs. Therefore, all boxes had the same appearance, a raised metallic box floor of approximately 5 cm above the true base, with ice packs mounted front and back.

Both the batteries and frozen ice packs had an effective life of around six hours and therefore were replaced each evening around 2.5 to 1.5 hours prior to sunset so that the treatments were running during the night. This period of the day was chosen as female blue tits usually roost overnight within their nest cavity prior to nest building and therefore should be present within the box to experience the temperature manipulation (Kluijver 1950; Perrins 1965a; Kidd et al. 2015). Temperatures were altered on average by ± 0.6 °C from the control boxes from sunset (around 6 pm) to midnight. Batteries and ice packs were replaced each evening; control boxes were also visited to achieve a uniform disturbance level across boxes. Some damage to equipment did occur (pecks to polystyrene inserts) and equipment was either repaired or replaced to ensure the treatments could continue. Experimental treatments could not be changed on two nights during the experiment (29^th^ March 2015 and 31^st^ March 2015) due to high winds. At the same time that treatments were changed several aspects of breeding phenology were recorded; signs of nest building (moss in box), signs of roosting (feathers and faeces), number of eggs, and whether the eggs were being incubated. From this we could calculate the nest building date (the date when moss first appeared in the nest box), clutch initiation date (the date on which the first egg was laid in a nest box) and clutch completion date (the date when the last egg was laid in a nest box).

In-nest temperatures were recorded for the entire duration of the experiment using iButton thermometers (DS1921G-F5, accurate to ± 1 °C, HomeChip Ltd) set to record temperature every 30 minutes. The iButtons were secured to the nest box wall using duct tape approximately 5 cm above the heating or cooling devices and half way between the front ice packs and the back ice packs. The effectiveness of the heating and cooling treatments during the experiment was assessed by calculating the mean temperature difference between treatments during the experimental periods (6 pm to 11:59 pm). The nights of the 29^th^ and 31^st^ March were excluded from these calculations because the experimental treatments did not run on those nights. Occupied nests were also excluded from the calculations after the first egg was laid, as the experimental treatments were halted at the appearance of the first egg. This type of experimental design was based on those previously employed on other nest box breeding passerine populations (Ardia et al. 2009; Pérez et al. 2008; YomTov & Wright 1993; Bryan & Bryant 1999; Vedder 2012; Alvarez & Barba 2014; Nager & van Noordwijk 1992). Previous manipulations have taken the form of heating and cooling usually during incubation (after laying has commenced) with heating devices attached to the outside of the nest box (Ardia et al. 2009; Pérez et al. 2008; YomTov & Wright 1993; Bryan & Bryant 1999) or within the nest box, below the nest itself (Vedder 2012; Alvarez & Barba 2014). Nager and Van Noordwijk (Nager & van Noordwijk 1992) heated and cooled nest boxes prior to laying through ice packs and heating devices located at the back of the nest box.

### Statistical Analyses

A power analysis was conducted prior to starting the experiment to determine whether differences were likely to be detected between treatments given the number of nest boxes available. Based on the average historical occupancy of the study boxes (67 %) we would expect an occupancy rate of 19 boxes per treatment. With this number the power analysis showed that we could expect a 75 % chance of detecting a mid to strong signal (an effect size of 0.4 or higher). We would expect a mid to high effect size based on previous work that has found a strong correlation between temperature and clutch initiation date (Perrins 1970; Perrins 1973). Furthermore, analyses of the relationship between spring temperature (mean of daily minimum and maximum temperatures for March and April) and clutch initiation date from 1960 to 2011 showed close to a 6 day delay in clutch initiation date for every 1 °C decline in mean temperature (EST = −5.71, SE = 0.08, P = <2e-16), therefore we would expect a strong signal from our experimental manipulation.

We performed statistical analyses to determine the influence of our experimental manipulations on breeding phenology and box occupancy. For the analyses of the influence of experimental treatments on box occupancy, we first tested if there was a difference in the number of boxes occupied between treatments. A chi-squared test of association was performed to determine if the number of occupied boxes differed significantly between treatments. It is also important to consider whether any other environmental factors could be influencing occupancy. Philopatry could have a role here with individuals or pairs returning to the same boxes every year. Across the length of this study 12 % of individuals (8 % of breeding attempts) returned to the same box they had previously bred in, suggesting that for the majority of individuals previous location is not the only driver of nest box choice. Therefore we used one way ANOVAs and a Welch’s T-Test to explore the difference in historical popularity, altitude and ambient temperature between occupied and unoccupied boxes. A Welch’s T-Test was used for local ambient temperature due to heteroscedasticity of temperature data. The whole dataset was used for these analyses as there was not enough data to split by treatment.

Analyses of the influence of experimental treatments on breeding phenology were performed using one way ANOVAs to determine if there are differences between phenology measures in each treatment group. The phenology measures considered here were the date of nest building (the date on which moss was first observed in the nest box), the clutch initiation date (date when the first egg is laid), and date of clutch completion (date when the last egg in the clutch is laid). Regression analyses were also performed to assess the influence of environmental variables on clutch initiation date. Environmental variables tested were altitude and local ambient mean temperature in a single linear regression to assess the combined influence of these variables on variance in phenology.

### Ethics statement

Work was subject to review by the Department of Zoology ethical committee, University of Oxford. All work adhered to UK standard requirements and was carried out under Natural England licence 20114732. This experimental methodology was discussed with, and approved by the Departmental Animal Welfare Ethical Review Body (AWERB). Field work took place in Wytham Woods (lat. 51°46’N, long. 1°20’W), private land that belongs to the University of Oxford; for permission contact the Conservator, Nigel Fisher. No endangered or protected species were involved in the study. A breakdown of nest building, abandonment and chick mortality by treatment can be found in supporting information Table S1, however these rates did not differ from those in non-experimental boxes.

## Results

### The influence of experimental treatment on occupancy

Occupancy in our experimental boxes was 16 %, which was a decline of 73 % from the preceding year (2014). Adjacent non-experimental boxes had a blue tit occupancy of 14 % (Table 1), a decrease of 48 % from 2014 occupancy. The number of great tits in these boxes also declined from 2014 to 2015, and to a greater extent, by 54 % (35 occupied in 2014 and 19 in 2015). Occupied boxes had higher historical popularity than unoccupied boxes (F(1,82) = 6.15, P = 0.02, see Figure 3), and this pattern was consistent across treatments. It should be noted that three of the occupied experimental boxes (N = 17) were occupied by marsh tits (*Poecile palustris).* As these are a different species to our study species, this could influence some of our results. Therefore we removed these nests from our analyses, leaving a sample size of N = 14.

**Figure 3:**
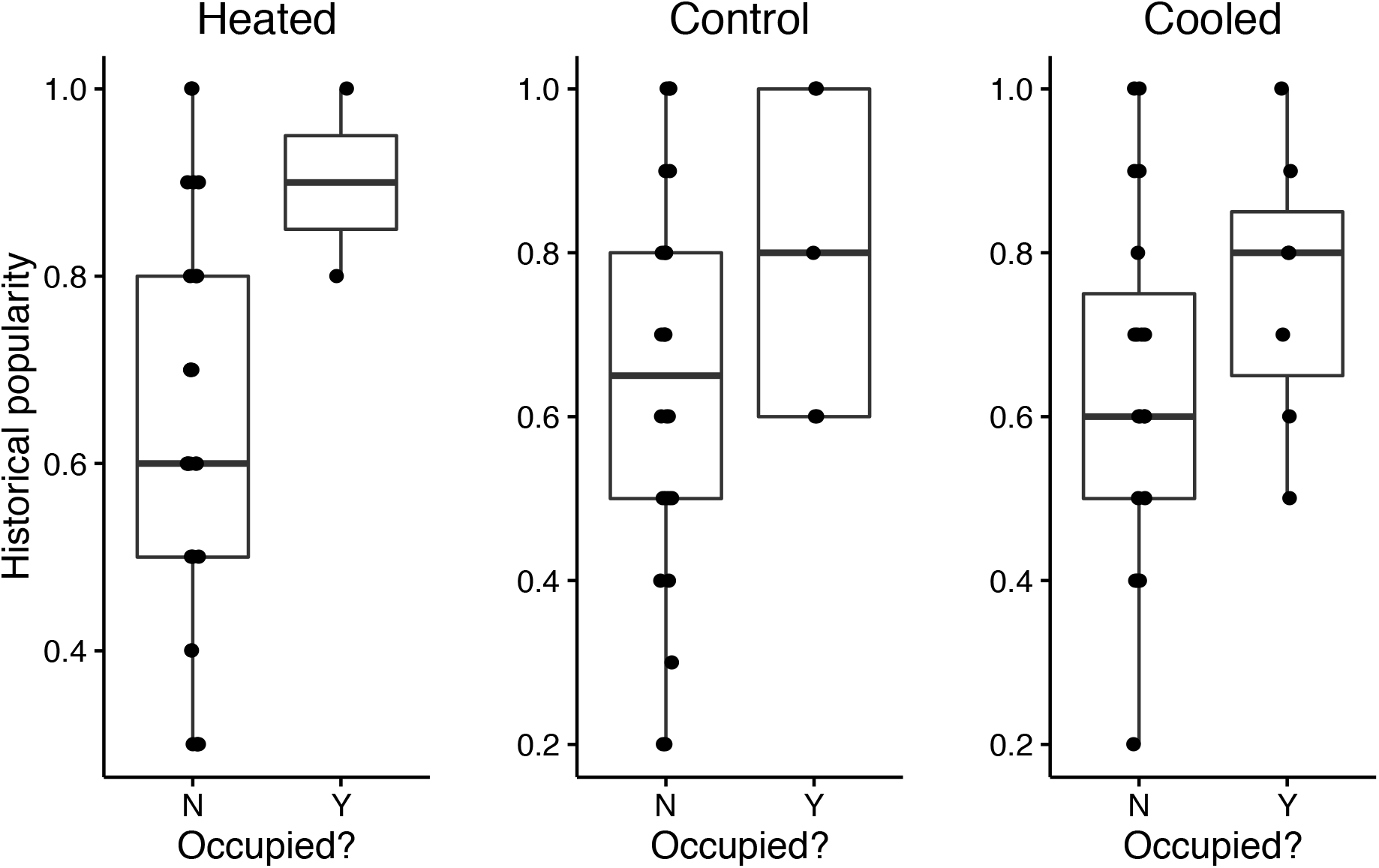
Boxplots of the historical popularity of occupied and unoccupied boxes by treatment. Raw data are plotted as points and the box plots show the median and interquartile and full range for each treatment. Historical popularity represents the proportion of years a box was occupied since its placement in the woodland.

**Table 1:** Numbers of occupied and empty boxes by treatment

\* Adjacent non-experimental boxes were defined as those boxes within 100 m of an experimental box.
| Treatment | Occupied boxes | Empty boxes (not occupied by blue tits) | Total |
| --- | --- | --- | --- |
| **Cooled** | 7 (24%) | 22 | 29 |
| **Control** | 5 (17%) | 24 | 29 |
| **Heated** | 2 (7%) | 27 | 29 |
| **Adjacent non-experimental \*** | 13 (14%) | 82 | 95 |

### Does in-nest temperature influence nest box choice?

The number of nest boxes occupied during our experimental period varied between treatments (Table 1). Cooled boxes had the highest proportion (24 %) of boxes occupied, followed by control boxes (17 %) and heated boxes (7 %) have the lowest levels of occupancy. These differences were not statistically significant (χ^2^ = 3.97, DF = 2, P = 0.14). We explored additional factors that may explain variation in occupancy. Neither altitude nor local mean ambient temperature differed significantly between occupied and unoccupied boxes (Altitude results, F(1,82) = 0.98, P = 0.33, local mean ambient temperature results - T = −1.00, P = 0.33, DF = 13.74, see supporting information, 0).

### Does in-nest temperature influence breeding phenology?

For these analyses N = 13 (two heated boxes, five control boxes and six cooled boxes), one nest was removed from analyses due to lack of a clutch initiation date as laying occurred after the conclusion of experimental treatments. On average, birds nesting in heated boxes started nest building, laying and completed their clutches earlier than birds nesting in the control or cooled boxes (Figure 4), however none of these differences were statistically significant (nest building F(2,11) = 1.20, P = 0.34, clutch initiation date F(2,11) = 0.91, P = 0.43, clutch completion date F(2,11) = 0.91, P = 0.43) (for full ANOVA tables see supporting information, 0). The influence of environmental variables, altitude (EST = −0.15, SE = 0.08, P = 0.07) and local ambient temperature (EST = −2.52, SE = 4.67, P = 0.60), were also tested but showed no statistically significant relationship with clutch initiation date, the focal phenological variable here.

**Figure 4:**
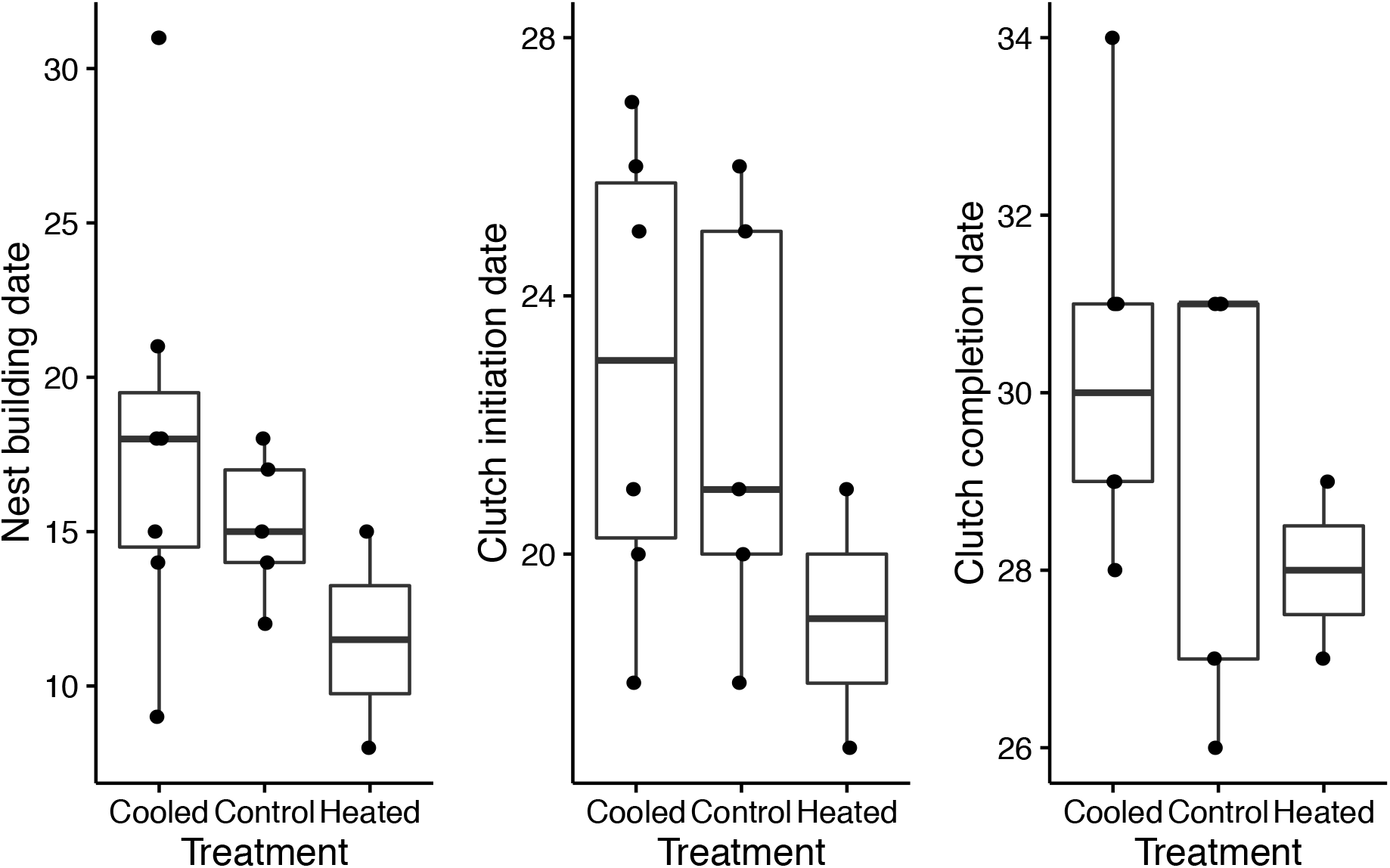
Boxplots of the dates of a) nest building, b) clutch initiation and c) clutch completion by treatment. Raw data are plotted as points and the box plots show the median and interquartile range for each treatment. Dates are all plotted in April days (April 1^st^ = 1)

## Discussion

### Experimental manipulation corresponded with low occupancy of nest boxes

This study experimentally tested whether nocturnal in-nest temperatures act as a cue for nest box choice and clutch initiation date in a population of wild blue tits. The experimental manipulation carried out in this study altered the minimum daily temperature experienced by individuals, either elevating it or reducing it dependent on treatment group. One of the defining findings of this study is a substantial and unforeseen decline in the occupancy of our experimental boxes to 16 %. This is considerably below the expected occupancy levels, which since 2008 have averaged 67 %. This 67 % occupancy would have given us approximately 19 boxes per treatment, which according to the preliminary power analysis conducted, would have been sufficient to detect the mid to high signal we expected (effect size of 0.4 or greater). However, due to circumstances beyond our control, this sample size was not achieved. The potential to increase our sample size was limited by the availability of blue tit only nest boxes. While more blue tit boxes are available in the woodland the remainder are located either in different habitat types or spatially distant from the study boxes. The inclusion of these other boxes would have introduced confounding factors and been logistically difficult to manage. Changes of equipment had to be achieved within a two hour window each evening. However, with the sample size available, we should have been able to detect a signal if one had been present and box occupancy was at the expected levels.

The low occupancy levels achieved in this experiment have resulted from several sources. Firstly, the number of nesting blue tit pairs declined across the whole of Wytham Woods in 2015, reducing by around 40 % (Table 2). Secondly, it was also clear that the parent blue tits found the disturbance and/or altering of the box appearance a deterrent from nesting in these boxes. Evidence that the birds were disturbed by the presence of the equipment was the pecking of polystyrene mounts in many of the boxes. This demonstrates the birds were not deterred from entering the box but also appeared to want to remove the heating and cooling devices. Evidence that birds entered the experimental boxes was also found in the form of signs of roost (faeces and feathers) found in several of the experimental boxes. However this use of boxes did not translate into nest building. Smaller nest cavities have been linked to higher nestling mortality (Mertens 1977) and could therefore deter parent birds. Additionally, the blue tit occupancy of the experimental nest boxes declined by a larger proportion than the non-experimental boxes (73 % and 48 % respectively, see Table 2). However, this cannot definitively be attributed to our experimental treatment because declines in both blue tit and great tit numbers were seen throughout the whole of Wytham Woods from 2014 to 2015. The lack of increased blue tit occupancy in the surrounding non-experimental boxes, suggests that birds deterred from the experimental boxes did not relocate to nearby unaltered boxes. However, disturbance did not seem to influence marsh tits to the same degree, their numbers remained at a similar level of occupancy to historical records, suggesting they were less deterred by the experimental set up than blue tits. Further refinements of techniques to minimise the amount of equipment required to heat and cool boxes would improve these experimental techniques. Caution should be employed for future blue tit nest box manipulations.

**Table 2:** Numbers of blue tit occupied nest boxes 2014 and 2015

|  | Boxes occupied 2014 | Boxes occupied 2015<br>(year of experiment) | Proportional<br>difference |
| --- | --- | --- | --- |
| **Whole woodland \*** | 430 | 258 | - 40% |
| **Non-experimental**<br>**boxes** | 378 | 244 | - 35% |
| **Experimental boxes** | 52 | 14** | - 73% |
| **Adjacent non-**<br>**experimental boxes** | 27 | 13 | - 48% |

### The impact of low sample size on statistical analyses

The sample size achieved in this study was exceptionally low, this heavily shaped our ability to draw statistically supported conclusions from this experimental work. With a sample size of 13 across all three treatments (two heated, five control, and six cooled), very little could be ascertained statistically from the resulting dataset. As analyses were planned prior to running of the experiment, they were duly conducted. However, even if statistically significant results are found, we cannot confidently reject a null hypothesis. The inability to determine statistical results stems from fundamental issues with multiple testing and p-values.

The rise of inappropriate and misleading statistical analyses is a growing concern across many scientific disciplines, in particular psychology, medicine, ecology, and evolution (Mcshane et al. 2017; Horton 2015; Parker et al. 2016; Forstmeier et al. 2016). It is something we must consider when analysing the data generated in this study and interpreting the results. A focus and reliance on p-values and null hypothesis testing, passing of a 0.05 threshold indicating a rejection of the null hypothesis, can be especially problematic for multiple comparisons on small datasets (Gelman & Loken 2014). When we accept a 5 % change of a Type 1 error (false positive result) over the course of multiple analyses, either on the same dataset or across studies, then 5 % of statistically significant results will actually have arisen if the null hypothesis were true (Forstmeier et al. 2016; Nuzzo 2014). Whether a result is deemed statistically significant or not is particularly changeable for small datasets where errors are high relative to the signal and the influence of individual data points is strong. Given this, there have been several calls to abandon p-values in statistics (Mcshane et al. 2017; Good 1988; Cumming 2014), although this call is not universal (Savalei & Dunn 2015).

Given the small sample size here, and the multiple comparisons conducted, the results should be analysed with caution. Multiple comparisons had to be conducted as there were multiple response variables to a focal explanatory variable (treatment), which were all chosen *a priori.* As a result, we move away from discussion of statistical significance in this study, as it cannot be relied upon in this instance (Mcshane et al. 2017). Instead we focus on the effect size of any trends found.

### Trends found

Trends shown in this analysis are that historical occupancy levels (popularity) also appear to predict occupancy during the experimental period. Boxes occupied during our experiment were occupied for, on average, 15.6 % more of the historical time period than the unoccupied boxes. It seems logical that the historically most popular boxes are also those occupied most readily during the experimental period. Neither altitude nor local mean temperature appear to explain popularity, either historically or during this experiment. Further analyses of different box positions would be required, manipulating specific aspects of the box environment, for example, aspect or oak density, to determine the drivers of occupancy/popularity of nesting sites. Although cooled boxes had the highest number of occupied boxes, this was statistically indistinguishable from control or heated. The difference between treatments was a maximum of four boxes. Past studies have found a slight preference for cooler nest boxes (Dhondt & Eyckerman 1979), however, this could also be a chance difference, substantially more analyses would be required to draw definitive conclusions.

No influence was found of in-nest temperature on breeding phenology. There was some overlap between the cooled and control groups for all measures of phenology, however, mean differences had effect sizes of several days. On average heated boxes initiated nest building and egg laying approximately three to four days prior to control boxes and six to four days prior to cooled boxes. The magnitude of these effect sizes from < 1°C change in in-nest temperature, predominantly driven by the heated treatment, suggest we cannot rule out an influence on nest box temperature on breeding phenology even if it could not be statistically supported in this study. No influence was found of environmental variables on clutch initiation dates. As ambient temperature typically correlates strongly with clutch initiation dates across our long-term data, we would expect that in this case the sample size was too low to distinguish a signal or the influence of ambient temperature was dampened by the experimental manipulation. However, from the current data, we cannot discriminate between the two possibilities. We were unable to find statistical support for the relationships observed by Dhondt and Eyckerman (Dhondt & Eyckerman 1979) and O’Connor (O’Connor 1978), similarly to Nager and Van Noordwijk (Nager & van Noordwijk 1992) we found little effect of in-nest manipulations on clutch initiation date. Due to the difficulty of wild experimental analyses and the low sample size achieved here, this result does not necessarily indicate a statistical null result. We would strongly encourage further replicated analyses of this type with higher sample sizes and across multiple years, before a robust conclusion can be reached.

## Acknowledgements

This study would not have been possible without the hard work of Tony Price and John Hogg, who assisted in the design and manufactured the heating and cooling devices for this experimental work. We would like to extend our gratitude to both Tony and John for all of the time they put in to this project. We are also grateful to all of the Wytham fieldworkers who collected population census data on the Wytham great tits. In particular, we would like to give a special mention to Keith McMahon, Stephen Lang, Koosje Lamers and Nico Alioravainen, who assisted with the changing of batteries and ice packs for this experiment come rain or shine. This work was supported by NERC grant NE/K006274/1 to Ben Sheldon.

## Supporting information

***Table of breakdown of nest building, abandonment and chick mortality by treatment***

**Table S1:** Table of breakdown of nest building, abandonment and chick mortality by treatment

\* These figures include great tit and marsh tit nesting attempts.
|  | Nest<br>building | Abandonment pre-<br>laying | Abandonment post-<br>laying | Chick<br>mortality |
| --- | --- | --- | --- | --- |
| **Cooled** | 7/27<br>(26%) | 1/7<br>(14%) | 0 | 2/61<br>(3%) |
| **Control** | 5/27<br>(19%) | 0 | 0 | 4/37<br>(11%) |
| **Heated** | 2/27<br>(7%) | 0 | 0 | 0 |
| **Non-**<br>**experimental** | 43/50*<br>(86%) | 9/43*<br>(21%) | 0 | 17/113<br>(15%) |

***Supporting tables: Does in-nest temperature influence nest box choice?***

**Table S2:** Full ANOVA table from analysis of occupancy as a function of historical popularity

| Response = historical popularity |  |  |  |  |  |
| --- | --- | --- | --- | --- | --- |
|  | DF | Sum of squares | Mean squared | F value | P |
| **Occupied** | 1 | 0.28 | 0.28 | 6.15 | 0.02 |
| **Residuals** | 82 | 3.78 | 0.05 |  |  |

**Table S3:** Full ANOVA table from analysis of occupancy as a function of altitude

| Response = altitude |  |  |  |  |  |
| --- | --- | --- | --- | --- | --- |
|  | DF | Sum of squares | Mean squared | F value | P |
| **Occupied** | 1 | 125.9 | 125.92 | 0.98 | 0.33 |
| **Residuals** | 82 | 10533.5 | 128.46 |  |  |

**Table S4:** Full table of Welch’s T-Test of occupancy as a function of mean temperature

| Response = mean temperature |  |
| --- | --- |
| **T** | -1 |
| **DF** | 13.74 |
| **P** | 0.33 |

***Supporting tables: Does in-nest temperature influence breeding phenology?***

**Table S5:**
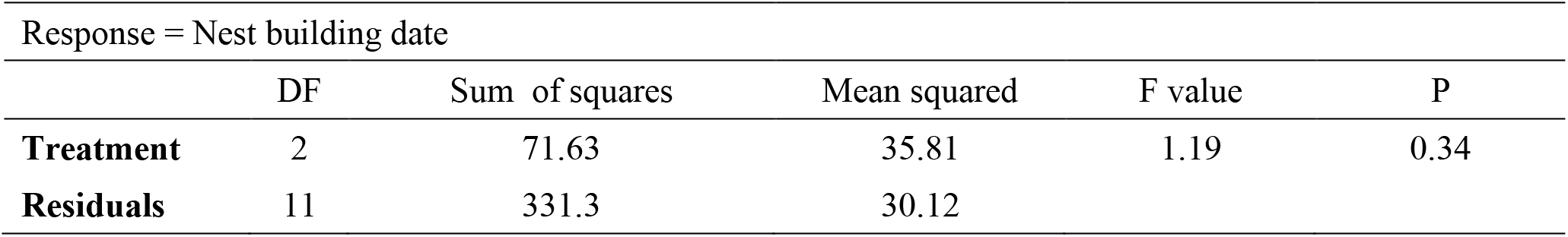
Full ANOVA table of nest building date as a function of treatment

**Table S6:** Full ANOVA table of lay date as a function of treatment

| Response = Lay date |  |  |  |  |  |
| --- | --- | --- | --- | --- | --- |
|  | DF | Sum of squares | Mean squared | F value | P |
| **Treatment** | 2 | 22.09 | 11.05 | 0.91 | 0.43 |
| **Residuals** | 11 | 120.83 | 12.08 |  |  |

**Table S7:** Full ANOVA table of clutch completion date as a function of treatment

| Response = Clutch completion date |  |  |  |  |  |
| --- | --- | --- | --- | --- | --- |
|  | DF | Sum of squares | Mean squared | F value | P |
| **Treatment** | 2 | 9.097 | 4.55 | 0.91 | 0.43 |
| **Residuals** | 11 | 50.13 | 5.01 |  |  |

**Table S8:**
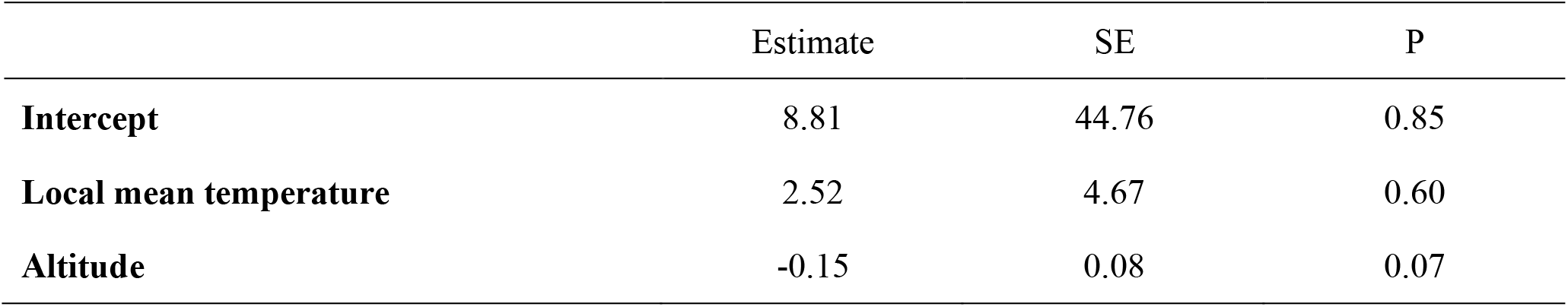
Table of outputs from phenology linear regression

|  | Estimate | SE | P |
| --- | --- | --- | --- |
| **Intercept** | 8.81 | 44.76 | 0.85 |
| **Local mean temperature** | 2.52 | 4.67 | 0.60 |
| **Altitude** | -0.15 | 0.08 | 0.07 |

